# Awake sharp-wave ripples coincide with rhythmic cortical synchronization

**DOI:** 10.64898/2026.09.29.754885

**Authors:** Tae Ho Lim, Xiaomin Zhang

## Abstract

Analysis of the Allen Institute Visual Coding Neuropixels dataset revealed a ∼5 Hz synchronization of visual cortical activity in awake, immobile mice. During bouts of this rhythm, synchronized cortical bursts coincided with changes in the occurrence of sharp-wave ripples (SWRs) and dentate spikes and in SWR properties. Cortical synchronization, SWRs, and dentate spikes were also coordinated with slower pupil-size fluctuation, revealing that awake SWRs are embedded within nested brainwide rhythms.

## Main Text

Low-frequency rhythms of the neocortex have been proposed to reflect a state of sensory disengagement during which neurons “beat in unison”.^1^ This notion dates back to the discovery of neural oscillations in 1929, when Hans Berger first reported prominent alpha waves (∼10 Hz) in the human electroencephalogram.^2^ Recent studies have found a similar ∼5 Hz rhythm to emerge in the mouse neocortex when disengaged from a task.^1^ Accumulating evidence suggests that this rhythm extends beyond the neocortex and into the thalamus, midbrain, and basal ganglia.^3–5^ Moreover, this rhythm appears to travel radially across sensory and motor cortices.^3,6^ Whether and how hippocampal activity is coordinated with this ∼5 Hz rhythm remains unclear.

Sharp-wave ripples (SWRs; 100–250 Hz) are prominent oscillatory events that emerge in the hippocampal CA1 during low-arousal states. They are most notably associated with memory consolidation, and during non-rapid eye movement sleep, they nest within slow waves and sleep spindles, forming a brainwide oscillatory architecture conducive to systems consolidation.^7^ How brainwide systems interact during awake SWRs remains much less understood, however. Dentate spikes (DSs) are large positive deflections of the dentate gyrus (DG) local field potential (LFP) that constitute another class of hippocampal population events. Although SWRs are generally considered distinct from DSs, a subset does temporally coincide,^8,9^ raising the question of the significance of this overlap.

To investigate how low-frequency cortical rhythms relate to hippocampal events in wakefulness, we analysed the Visual Coding dataset made publicly available by the Allen Brain Institute.^10,11^ In 40 mice, 6 Neuropixels probes each targeted a different visual cortical (vCtx) area (V1, RL, AL, L, PM, AM) while also sampling structures including the visual thalamus, midbrain, and hippocampus (Fig. 1a; Extended Data Fig. 1). Running speed and pupil diameter were recorded concurrently. vCtx multi-unit activity (MUA) revealed bouts in which baseline activity dropped markedly while rhythmic low-frequency (3–10 Hz) activity emerged (Fig. 1b). We refer to individual rhythmic bursts within these bouts as synchronization events.

**Fig. 1:**
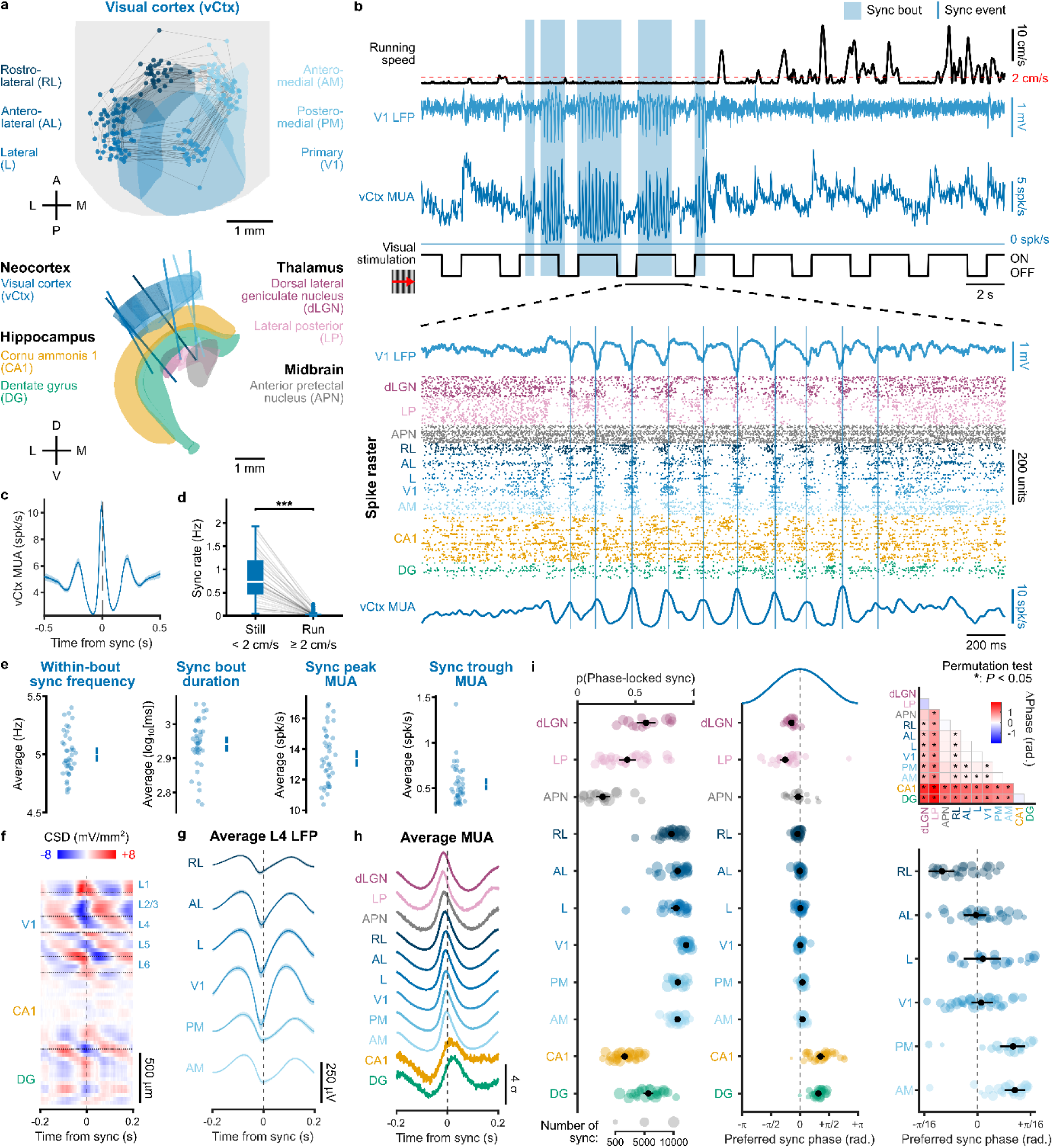
The visual cortex synchronizes at ∼5 Hz during immobility. **a**, Recording geometry and targeted regions.^28^ **b**, Example behavioural and electrophysiological recordings during drifting gratings stimulation. **c**, vCtx MUA aligned to synchronization (*n* = 40 mice; 189,152 events). Line and shading indicate mean and 95% hierarchical-bootstrap CI. **d**, Synchronization rate during immobility and movement (*n* = 40 mice; *P* = 3.57 × 10^-8^). **e**, Within-bout synchronization frequency, bout duration, and peak and trough vCtx MUA (*n* = 40 mice). Bars indicate 95% hierarchical-bootstrap CI. **f**, Example synchronization-aligned cortico-hippocampal CSD. **g**, Synchronization-aligned layer 4 LFP across vCtx areas (*n* = 19–33 mice). Line and shading indicate mean and 95% hierarchical-bootstrap CI. **h**, Mean normalized regional MUA aligned to synchronization (*n* = 14–36 mice). **i**, Regional fraction of phase-locked synchronization cycles and preferred synchronization phase (*n* = 14–36 mice). Pairwise regional permutation tests were Holm-corrected separately for locking fraction and phase difference. \**P* < 0.05.

Synchronization occurred more frequently during immobility than during movement (median, 0.731 versus 0.0234 Hz; *n* = 40 mice; *P* = 3.57 × 10^-8^; Fig. 1d). Within bouts, synchronization recurred at 5.00 Hz (95% CI, 4.94–5.06 Hz), with bouts lasting a geometric mean duration of 0.87 s (95% CI, 0.83–0.91 s). This rhythmic population activity alternated between peaks of 13.38 spk/s (95% CI, 12.85–13.94 spk/s) and troughs of 0.538 spk/s (95% CI, 0.478–0.608 spk/s; Fig. 1e). Although synchronization also emerged spontaneously, visual input strongly modulated its occurrence, consistent with previous reports (Extended Data Figs. 2, 3).^1,3,4,12–16^

Synchronization was accompanied by structured activity across cortical and subcortical regions. Synchronization-aligned current-source density (CSD) revealed a cortical layer 4 sink followed by a DG sink (Fig. 1f; Extended Data Fig. 2e). Regional MUA showed distinct synchronization phase preferences (Fig. 1i). Visual thalamic nuclei preferred to fire earlier in the cycle (dLGN: −0.462 rad.; LP: −0.810 rad.), whereas hippocampal areas preferred to fire later (CA1: 1.12 rad.; DG: 0.999 rad.). Although vCtx areas clustered in their phase preference, their systematic spatial ordering was consistent with previous reports of thalamocortical^5,13^ and radial propagation of synchronization.^3,6^

Could the delayed hippocampal phase preference reflect the coordination of hippocampal population events with cortical synchronization? To investigate this, we detected SWRs and DSs in the CA1 and DG LFP, respectively (Extended Data Figs. 4, 5). DSs were further classified as DS1 and DS2 depending on whether their dominant CSD sink was located in the outer (OML) or middle molecular layer (MML). SWR, DS1, and DS2 exhibited their largest sink probability in the stratum radiatum (SR), OML, and MML, respectively. Notably, synchronization was also associated with an increased MML sink probability, revealing a laminar signature shared with DS2 (Fig. 2e).

**Fig. 2:**
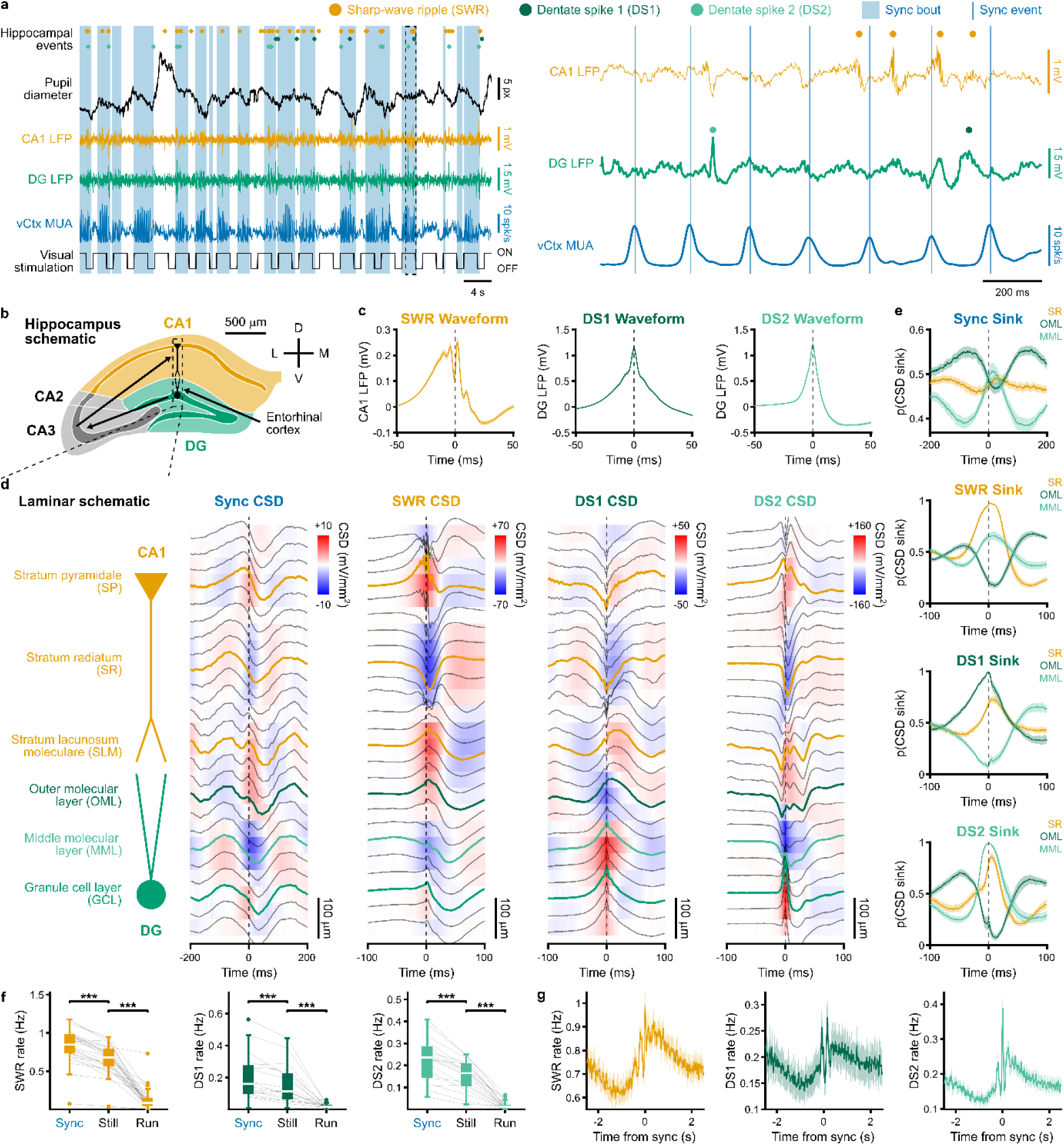
Hippocampal events coincide with cortical synchronization. **a**, Example recording during drifting gratings. **b**, Schematic adapted from the Allen Mouse Brain Atlas.^29^ **c**, SWR, DS1, and DS2 LFP waveforms (SWR, *n* = 33 mice, 321,445 events; DS1, *n* = 29, 40,606; DS2, *n* = 29, 36,379). **d**, CSD and LFP aligned to synchronization, SWR, DS1, and DS2. **e**, Negative CSD probability in SR, OML, and MML (*n* = 19 mice). In **c**, **e**, line/ shading denote mean/ 95% hierarchical-bootstrap CI. **f**, SWR/ DS1/ DS2 rates during synchronization bouts, immobility, and movement. Synchronization versus immobility: *n* = 26/ 18/ 18 mice, *P* = 4.98 × 10^-5^/ 7.86 × 10^-4^/ 7.86 × 10^-4^; immobility versus movement: *n* = 18/ 26/ 18 mice, *P* = 8.63 × 10^-4^/ 4.98 × 10^-5^/ 7.86 × 10^-4^. **g**, Synchronization-aligned rates (*n* = 26/ 18/ 18 mice); mean ± s.e.m. *P* values were Holm-corrected. \**P* < 0.05, *\*\*P* < 0.01, \*\*\**P* < 0.001.

Median SWR, DS1, and DS2 rates were all elevated during synchronization bouts compared with immobility overall (SWR, 0.847 versus 0.682 Hz, *n* = 26 mice, *P* = 4.98 × 10^-5^; DS1, 0.236 versus 0.165 Hz, *n* = 18 mice, *P* = 7.86 × 10^-4^; DS2, 0.114 versus 0.017 Hz, *n* = 18 mice, *P* = 7.86 × 10^-4^; Fig. 2f). Aligning hippocampal event rates to synchronization revealed a shared temporal structure, with a ∼5 Hz modulation nested within a slower, seconds-long modulation of event occurrence (Fig. 2g). This suggested that hippocampal events were organized across multiple temporal scales.

We therefore asked whether the slow component was related to slow pupil size dynamics (0.1–1 Hz), which track fluctuations in arousal.^15,17^ Synchronization, SWRs, DS1s, and DS2s all preferentially locked to the constriction phase of the pupil cycle (Fig. 3a; Extended Data Fig. 6). While this shared slow modulation could contribute to the enrichment of hippocampal events during synchronization bouts, SWRs, DS1s, and DS2s also phase-locked to the ∼5 Hz synchronization cycle (Fig. 3b). DS1 preferentially occurred near the synchronization trough (−1.91 rad.), whereas SWR (0.941 rad.) and DS2 (0.115 rad.) occurred preferentially near the synchronization peak. Hippocampal events were therefore organized across two slower brainwide timescales, reminiscent of the nested oscillatory architecture of sleep.

**Fig. 3:**
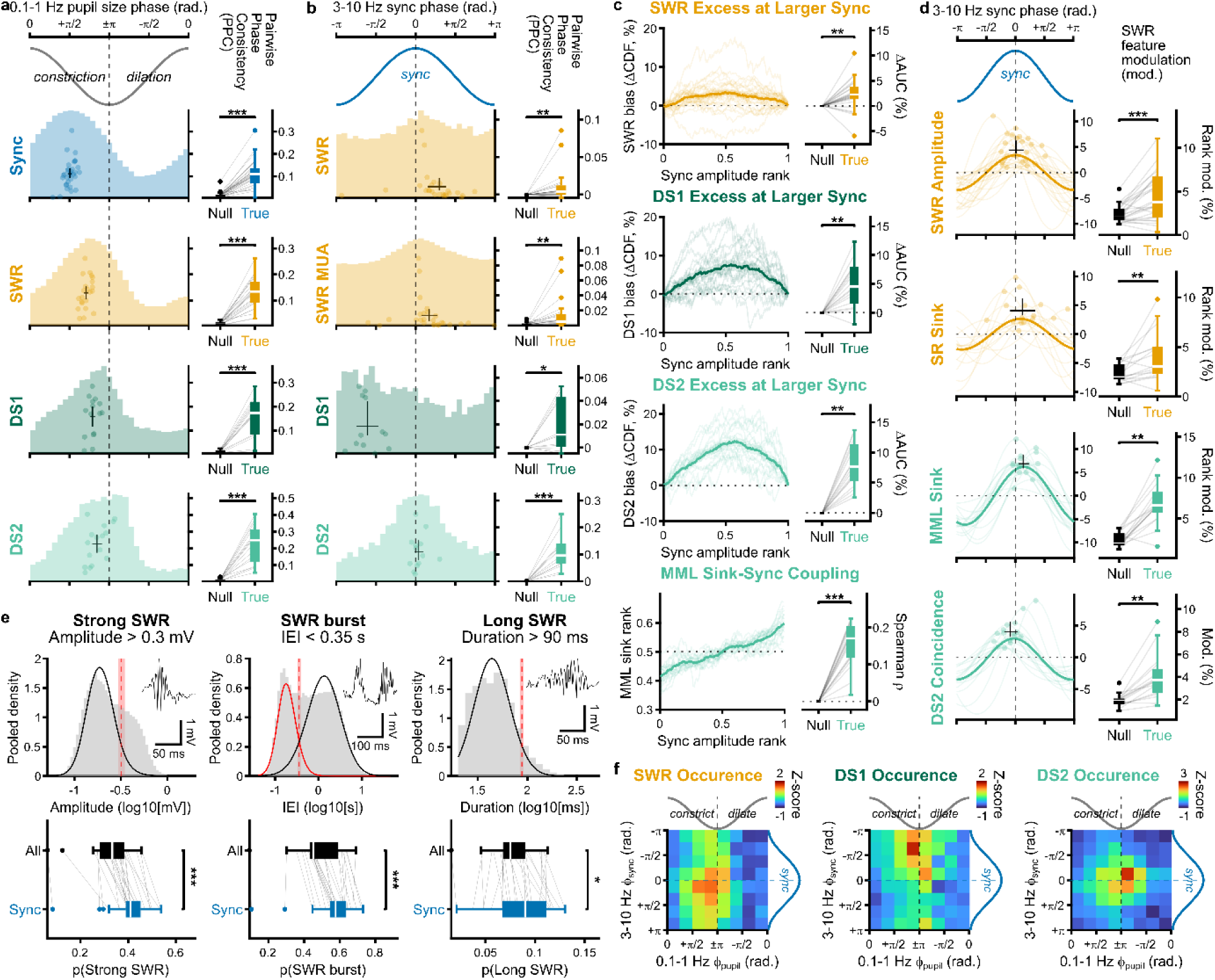
Cortical synchronization structures hippocampal events. **a**, Locking of synchronization, SWR, DS1, and DS2 to 0.1–1 Hz pupil phase during immobility. For synchronization/ SWR/ DS1/ DS2, *n* = 32/ 22/ 15/ 16 mice and *P* = 3.5 × 10^-6^/ 1.2 × 10^-4^/ 1.22 × 10^-4^/ 4.38 × 10^-4^. **b**, Locking of SWR, CA1 spikes occurring during SWRs, DS1, and DS2 to synchronization phase. Respectively, *n* = 25/ 26/ 12/ 13 mice and *P* = 0.002/ 0.005/ 0.016/ 9.77 × 10^-4^. **c**, Bias of hippocampal event-associated synchronization toward larger synchronization amplitude ranks and coupling of synchronization amplitude rank with associated MML sink strength rank (ΔAUC = AUC − 50%). For SWR/ DS1/ DS2, *n* = 26/ 17/ 16 mice, median ΔAUC = 2.26/ 4.54/ 7.67%, and *P* = 0.001/ 0.001/ 0.001; MML coupling, *n* = 19 mice, median Spearman ρ = 0.171; *P* = 5.27 × 10^-4^. **d**, First-harmonic modulation across synchronization phase of SWR amplitude rank, SR- and MML-sink-strength ranks, and SWR–DS2 coincidence. *n* = 26/ 16/ 16/ 16 mice and *P* = 9.7 × 10^-4^/ 0.005/ 0.002/ 0.002. **e**, Fitted cohort thresholds for strong, burst, and long SWRs were 0.320 mV (95% hierarchical-bootstrap CI, 0.294–0.349 mV), 0.356 s (0.322–0.400 s), and 88.13 ms (84.21–92.03 ms) (*n* = 25 mice), with applied cutoffs of 0.3 mV, 0.35 s, and 90 ms, respectively. These SWRs were enriched among synchronization-associated SWRs (*n* = 26 mice; *P* = 2.49 × 10^-5^/ 2.65 × 10^-5^/ 0.011). **f**, Descriptive joint pupil-synchronization phase within-mouse z-scored SWR, DS1, and DS2 event rates (*n* = 16–22 mice). Crosses in **a**, **b**, **d** indicate cohort phase and pairwise-phase consistency (PPC) or modulation with 95% hierarchical-bootstrap CI. *P* values were Holm-corrected within each comparison family. \**P* < 0.05, \*\**P* < 0.01, \*\*\**P* < 0.001.

Beyond temporal coincidence, we next asked whether hippocampal events were more likely to coincide with larger synchronization events. Indeed, coincidences with SWR, DS1, or DS2 were biased towards larger synchronization amplitudes (median ΔAUC = 2.26%, 4.54%, 7.67%, respectively; *P* = 0.001 for each; Fig. 3c). Interestingly, this bias and phase locking were most pronounced for DS2 (Extended Data Fig. 7). Consistent with this preferential relationship, synchronization amplitude covaried with MML sink strength (median Spearman ρ = 0.171; *n* = 19 mice; *P* = 5.27 × 10^-4^), more strongly than with the other hippocampal laminar sinks (Extended Data Fig. 7). This differential coupling may therefore reflect a circuit route through which cortical synchronization interacts with the hippocampus.

The relatively weak coupling of SWR occurrence to synchronization raised the possibility that SWR features may nevertheless vary across the synchronization cycle. Among the features examined, four showed significant phase modulation (Fig. 3d; Extended Data Fig. 7). SWR amplitude (preferred phase, 0.033 rad.) and SR sink strength (0.378 rad.) were both greatest near the synchronization peak, consistent with stronger ripples accompanying larger sharp waves. Interestingly, MML sink strength (0.422 rad.) and SWR–DS2 coincidence (−0.269 rad.) were similarly enhanced near the synchronization peak. Together, these observations raise the possibility that SWRs occurring near the peak of cortical synchronization may coincide with a distinctive convergence of CA3–CA1 and entorhinal–dentate input.

If stronger SWRs preferentially coincide with the synchronization peak, could synchronization-associated SWRs differ systematically from the broader SWR population? We found that synchronization-associated SWRs were biased towards larger amplitudes, lower frequencies, longer durations, shorter inter-event intervals (IEI), and stronger hippocampal laminar sinks (Extended Data Fig. 7). This prompted us to ask whether SWRs associated with extended replay were particularly enriched during synchronization bouts.^18,19^ We classified SWRs as strong (> 0.3 mV amplitude), burst (< 0.35 s IEI), or long (> 90 ms duration). Indeed, all three were significantly enriched among synchronization-associated SWRs (strong, median proportions, 0.410 versus 0.333, *P* = 2.49 × 10^-5^; burst, 0.576 versus 0.452, *P* = 2.65 × 10^-5^; long, 0.0901 versus 0.075, *P* = 0.011; Fig. 3e). Together, these findings reveal cortical synchronization as a previously unrecognized temporal context for a distinctive subset of awake SWRs.

In conclusion, our findings reveal how cortical synchronization is coordinated with hippocampal activity. Awake hippocampal events are nested within a slow 0.1–1 Hz pupil-associated rhythm and a faster ∼5 Hz cortical synchronization rhythm. This coordination extends beyond timing: larger synchronization events more often coincide with hippocampal events, while stronger SWRs occur near the synchronization peak. The close coupling between synchronization and MML activity implicates the entorhinal–dentate circuit in this coordination. Given the propagation of synchronization between the visual and retrosplenial cortices,^1,3,13,14^ a visual–retrosplenial–entorhinal circuit could provide one pathway by which cortical synchronization may reach the hippocampal formation.

The preferential coincidence of SWRs and DS2 during cortical synchronization is particularly intriguing because they have been proposed to reflect complementary “offline” and “online” modes of hippocampal processing, respectively.^8^ The tight coupling of DS2 to synchronization further raises the possibility that the brainwide activation previously associated with DS2 partly reflects this cortical rhythm. The coincidence of SWRs with such brainwide activation, however, is less readily explained by their classical “offline” role. Yet recent human and primate studies increasingly implicate a subset of awake SWRs in ongoing perceptual and cognitive processes.^20–25^ Synchronization-associated SWRs may therefore reflect a poorly understood subset of awake SWR activity.

More broadly, the nesting of awake SWRs within slow pupil size fluctuations and faster synchronization rhythms echoes the nesting of sleep SWRs within slow waves and sleep spindles. This is not to say that these architectures are equivalent; rather, they may reflect a common organizing principle of SWR-associated cortico-hippocampal communication. In sleep, slow waves preferentially propagate along anatomically structured cortical pathways,^26^ while in wakefulness, low-frequency cortical waves are likewise constrained by axonal architecture.^3^ Slow pupil size fluctuations in wakefulness^17^ and slow waves in sleep^27^ are also closely linked to changes in neuromodulatory tone. Anatomical connectivity may therefore constrain where brainwide synchronization propagates, while neuromodulatory state shapes when it emerges, together defining windows for cortico-hippocampal communication.

## Methods

### Dataset

We analysed the publicly available Allen Brain Observatory Visual Coding Neuropixels dataset.^10,11^ The dataset consisted of recordings from 58 awake, head-fixed mice during passive visual stimulation. Up to 6 Neuropixels probes were recorded simultaneously, each targeting one of six visual cortical areas (V1, RL, AL, L, PM, and AM). Electrophysiological data included local field potentials (LFP) and spike-sorted unit activity, and behavioural measurements included running velocity and pupil size. The primary cohort consisted of mice with all 6 Neuropixels probes inserted, each with valid LFP recordings (*n* = 40 mice), ensuring consistent cortical coverage across mice. Individual analyses used subsets of this cohort depending on data availability, anatomical sampling, and analysis-specific criteria. Allen-provided structure annotations were used to assign recording channels and units to brain regions. Visual cortical layers were assigned by mapping channel Common Coordinate Framework (CCF) coordinates^30^ onto a 10 μm modified Allen CCF annotation volume^28^ using cortex-lab/AllenCCF utilities (https://github.com/cortex-lab/allenCCF). Probe trajectories and insertion sites were reconstructed from supplied CCF channel coordinates.

### Preprocessing

Analysis was restricted to the common recording period with valid data from all 6 probes. Allen-designated invalid intervals and, where present, data from the first optogenetic stimulation onward were excluded. Spike-sorted units were classified as good if their inter-spike interval violation < 0.5, presence ratio > 0.9, and amplitude cutoff < 0.1. Only good units were retained for subsequent analyses. Visual cortical units were further classified by laminar location as supragranular (SG; layers 1 and 2/3), layer 4, or infragranular (IG; layers 5–6), and by their trough-to-peak waveform duration as putative fast-spiking (FS; < 0.425 ms) or regular-spiking (RS; ≥ 0.425 ms).

Running velocity was linearly interpolated to 60 Hz and smoothed with a Gaussian kernel (σ = 35 ms). Unless otherwise stated, Gaussian smoothing kernels were L1-normalized. Running velocity was rectified to obtain the running speed, and immobility was defined as periods in which running speed < 2 cm/s lasting ≥ 3 s, with remaining periods considered movement.

Pupil diameter was calculated as the mean of pupil width and height and linearly interpolated to 30 Hz. To capture pupil size fluctuations, pupil diameter was band-pass filtered at 0.1–1 Hz using a third-order zero-phase Butterworth filter. Instantaneous phase was obtained from the analytic signal using the Hilbert transform, such that 0 rad. corresponds to maximal pupil size, +π/2 rad. to constriction, ±π rad. to minimal pupil size, and −π/2 rad. to dilation. Pupil-phase analyses were restricted to valid immobility periods.

### Data analysis

#### Cortical synchronization

Spikes from good vCtx units were binned at 1 ms resolution and averaged across units to obtain vCtx multi-unit activity (MUA). MUA was band-pass filtered at 3–10 Hz using a fourth-order zero-phase Butterworth filter, and instantaneous phase and amplitude were obtained by Hilbert transform. Baseline MUA was estimated as the moving 20^th^ percentile of MUA over 300 ms of MUA smoothed with a Gaussian kernel (σ = 10 ms). The 3–10 Hz amplitude and baseline MUA were smoothed with a Gaussian kernel (σ = 125 ms).

Synchronization bouts were defined as periods with 3–10 Hz amplitude > 2 spk/s and baseline MUA < 4 spk/s. Candidate bouts were retained when maximum 3–10 Hz amplitude > 3 spk/s, minimum baseline MUA < 3 spk/s, and the unwrapped 3–10 Hz phase progression spanned at least two oscillatory cycles. Within-bout synchronization frequency was calculated as the total unwrapped 3–10 Hz phase progression, expressed in cycles, divided by bout duration.

Individual synchronization events were defined as peaks of the 3–10 Hz oscillation occurring within synchronization bouts. Phase was linearly interpolated between successive synchronization peaks within a bout to yield synchronization phase. Thus, synchronization occurred at 0 rad., and the surrounding ±π phase boundaries delineated individual synchronization cycles. Synchronization amplitude was measured from the vCtx MUA smoothed with a Gaussian kernel (σ = 10 ms). Peak MUA was defined as the maximum instantaneous MUA within each cycle and trough MUA as the minimum instantaneous MUA between consecutive synchronization peaks within the same bout.

Synchronization-aligned visual cortical LFP analyses required ≥ 100 synchronization events per mouse and area, and regional MUA analyses additionally required ≥ 20 good units per mouse and region. Regional phase-locking analyses required ≥ 20 good units and ≥ 200 eligible synchronization cycles per mouse and region, with eligible cycles containing ≥ 10 regional spikes. Pairwise phase consistency (PPC) and mean spike phase were calculated for each eligible cycle, and cycles with PPC ≥ 0.04 were classified as phase-locked. Regional preferred phase required ≥ 20 phase-locked cycles. Unit-level phase-locking analyses required ≥ 20 synchronization-associated spikes per unit, with PPC ≥ 0.04 defining phase locking. Mouse-level summaries required ≥ 5 eligible units, and preferred-phase summaries required ≥ 5 phase-locked units.

#### Sharp-wave ripples

Sharp-wave ripples (SWRs) were detected on probes containing at least one CA1 channel. For eligible probes, the stratum pyramidale (SP) channel was assigned to the CA1 channel with the highest ripple score. Ripple score was calculated as the ratio of 120–200 Hz to 70–300 Hz power estimated using Welch’s method.

SP LFP was band-pass filtered at 100–250 Hz using a fourth-order zero-phase Butterworth filter to obtain the ripple LFP. Ripple amplitude was obtained from the magnitude of the analytic signal and smoothed with a Gaussian kernel (σ = 4 ms) to obtain the ripple envelope. Candidate SWR windows were defined when envelope exceeded 6 times the median absolute ripple LFP and were retained if the maximum ripple envelope exceeded 9 times this value.

Candidate SWRs separated by < 25 ms were merged and were retained if their duration was 20–350 ms and contained at least three ripple cycles. SWR peak was defined as the time of maximum ripple amplitude within the SWR window. SWR amplitude was defined as the corresponding amplitude, and SWR frequency was calculated as the unwrapped 100–250 Hz phase progression, expressed in cycles, divided by the SWR duration.

SWR detection results were visually inspected for each probe; SP channels could be reassigned and probes excluded. Mice with at least four manually validated SWR probes spanning the dorsal-to-intermediate CA1 were retained for SWR-related analyses, ensuring consistent SWR coverage. Subsequently, SWRs separated by ≤ 30 ms were merged across probes. Merged SWR peak, amplitude, and frequency were defined by the constituent SWR with the largest amplitude, whereas duration was calculated from the merged SWR window. SWR inter-event interval (IEI) was calculated between consecutive merged SWR peaks.

#### Dentate spikes and current-source density

Dentate spikes (DS) were detected on the four dorsal-most probes (A, B, C, F) with consecutive CA1–DG channels. DG LFP was band-pass filtered at 1–200 Hz using a fourth-order zero-phase Butterworth filter, and positive excursions 7 times the median absolute filtered LFP were detected. DS time and amplitude were defined by the maximal filtered LFP within the excursion. DSs separated by < 50 ms were merged retaining the constituent DS with the largest amplitude. The DG channel with the greatest number of detected DS was assigned the granule cell layer (GCL), and subsequent DS analysis used only DSs detected on this channel.

Current-source density (CSD) was calculated as the negative second spatial derivative of the LFP. Adjacent channels were separated by 40 μm along the probe axis, and CSD was spatially smoothed with a Gaussian kernel (σ = 50 μm). Negative CSD values hence reflect current sinks. Hippocampal laminae were identified physiologically. The stratum radiatum (SR) was assigned to the channel below SP with the largest SWR-aligned CSD sink. To identify the stratum lacunosum-moleculare (SLM), theta epochs were detected when 5–12 Hz DG LFP power, estimated using a multitaper spectrogram implemented in Chronux 2.12^31^ (4 s window, 2 s step; time-bandwidth product, 3; 5 tapers) and smoothed across 10 time bins, exceeded its 90^th^ percentile. Theta peaks were detected in the 5–12 Hz SP LFP during theta epochs, and the SLM was assigned to the channel with the greatest theta-peak-aligned sink between the SR and GCL.

To identify the outer molecular layer (OML) and middle molecular layer (MML), the SLM–GCL CSD profile of each DS was z-scored then subjected to principal component analysis. Mean CSD profiles from the extreme positive and negative halves of the PC1 distribution revealed two characteristic sink locations, one closer to the SLM and the other closer to the GCL. They were assigned as the OML and the MML channels, respectively. DSs with a larger OML than MML sink were classified as DS1, whereas DSs with a larger MML than OML sink were classified as DS2.

DS detection and laminar assignments were visually inspected, and probes could be excluded or GCL, SLM, OML, and MML channels could be manually reassigned. Mice with at least two eligible probes were retained for DS-related analyses. DS separated by ≤ 100 ms were subsequently merged across probes, with the constituent DS of the largest amplitude defining the merged DS time, amplitude, and DS1/2 classification. For hippocampal CSD analyses, one manually selected probe with a well-resolved hippocampal CSD profile was used for each mouse; probe-matched SWR and DS events, rather than merged events, were used in these analyses.

For event-aligned CSD analyses, sink probability was quantified as the fraction of aligned events with negative CSD at each time point. For sink-strength analyses, CSD was low-pass filtered at 30 Hz, and sink strength was defined as the magnitude of the most negative CSD within the corresponding event window. For analyses of discrete CSD sinks, contiguous periods of negative filtered CSD were identified as individual sinks, with sink time and strength defined by the time and magnitude of the minimum CSD within each period.

#### Event and phase analyses

SWR, DS1, and DS2 rates were calculated as event counts divided by valid recording duration within synchronization bouts, immobility, or movement. Synchronization bouts remained part of the overall immobility condition. Peri-synchronization event rates were calculated in 1 ms bins and smoothed with a Gaussian kernel (σ = 10 ms).

Locking to slow pupil-size dynamics was quantified by sampling 0.1–1 Hz pupil phase at synchronization, SWR, DS1, and DS2 times during immobility. Mice required ≥ 100 events of the relevant class. Preferred phase and PPC were calculated for each mouse. Null PPC was estimated from 100,000 non-zero circular shifts of event positions through the concatenated valid immobility samples. Frequency specificity of synchronization locking was analysed analogously after filtering pupil diameter at 0.01–0.1, 0.01–1, 0.01–10, 0.1–10, and 1–10 Hz.

To test whether event occurrence was associated with pupil size independently of phase, complete pupil cycles were identified, and pupil diameter was interpolated as a function of phase within each cycle. Pupil size at each event’s phase was ranked across complete cycles, yielding a phase-conditioned pupil-size rank. Null ranks were generated by non-zero circular shifts of pupil-cycle identity while retaining event phase.

Locking to synchronization phase was quantified for SWRs, good CA1 unit spikes occurring during SWRs, DS1, and DS2. Mice required ≥ 100 observations of the relevant class. Preferred phase and PPC were calculated for each mouse. Null PPC was estimated from 100,000 random circular temporal shifts within synchronization cycles, with one shift shared by all observations occurring in the same cycle.

Phase-dependent modulation of SWR features and event-associated CSD sink strengths was quantified using first-harmonic models. Continuous outcomes were converted to within-mouse quantile ranks and fitted with sine and cosine terms; binary outcomes were fitted using logistic first-harmonic models. Analyses required ≥ 50 SWRs, with binary analyses additionally requiring ≥ 10 observations in each class. Modulation was compared with a null generated from 10,000 within-mouse permutations of outcomes across observed phases. Continuously sampled vCtx MUA and 3–10 Hz MUA amplitude were fitted across pupil phase without rank transformation and compared with circular-shift nulls. Discrete CSD sink strengths were ranked within mouse and analysed using the corresponding first-harmonic permutation procedure.

Joint pupil-synchronization phase maps were calculated by dividing both phases into 8 bins. Within each mouse, SWR, DS1, and DS2 rates in each of the resulting 8 × 8 bins were calculated as event count divided by occupancy duration, z-scored across bins and then averaged across mice.

#### Feature analyses

To determine whether hippocampal events preferentially occurred during larger synchronization events, synchronization cycles were ranked within mouse by peak vCtx MUA. Cycles containing ≥ 1 SWR, DS1, or DS2 were compared with event-free cycles using the Mann-Whitney AUC, equivalent to the probability that an event-containing cycle had greater synchronization amplitude than an event-free cycle.

Analyses required ≥ 50 event-containing and ≥ 2 event-free synchronization cycles per mouse. ΔAUC was defined as AUC − 50%, such that positive values indicate preferential association with larger synchronization events. Synchronization amplitude and MML sink strength were additionally related using Spearman correlation; these analyses required ≥ 10 eligible synchronization cycles per mouse.

Biases in synchronization-associated SWR properties were quantified analogously by comparing synchronization-associated with non-synchronization-associated SWRs. SWR amplitude, frequency, duration, shorter IEI, and SR, SLM, OML, and MML sink strengths were ranked within mouse, and their association with synchronization was summarized by ΔAUC. Analyses required ≥ 50 synchronization-associated and ≥ 2 non-synchronization-associated SWRs.

SWRs were additionally classified as strong (> 0.3 mV amplitude), burst, or long (> 90 ms duration). Consecutive SWRs separated by < 0.35 s were grouped, and SWRs belonging to groups containing ≥ 2 events were classified as burst SWRs. These fixed thresholds were selected to approximate cohort-derived thresholds. For threshold estimation, mice required > 1,000 eligible observations. Ripple amplitude and duration were analysed after log10 transformations: the strong threshold was estimated as the upper 95^th^ percentile of a Gaussian distribution fitted from the lower 20% of amplitudes, and the long threshold from the upper 95^th^ percentile of a Gaussian whose spread was estimated from the lower half of the duration distribution. The burst threshold was estimated as the crossing of a two-component Gaussian mixture fitted to log10 IEIs between consecutive SWRs occurring within the same immobility period. Animal thresholds were averaged in log space and 95% confidence intervals estimated by hierarchical bootstrap, yielding cohort thresholds of 0.320 mV, 0.356 s, and 88.13 ms, respectively. Enrichment was quantified by comparing the proportion of each SWR class among synchronization-associated SWRs with its proportion among all SWRs.

#### Visual stimulus analyses

Visual stimulus rate analyses were restricted to valid immobility. Synchronization, SWR, DS1, and DS2 rates were calculated during Gabors, full-field flashes, drifting gratings, contrast gratings, static gratings, natural scenes, natural movies 1 and 3, dot motion, and spontaneous activity as event counts divided by the corresponding immobile duration. Spontaneous presentations lasting > 10 s were retained for the spontaneous reference.

For stimulus-aligned analyses, synchronization-bout occurrence and pupil phase were aligned to stimulus onset using valid immobile samples. Natural movie motion energy was calculated at 30 Hz as the root-mean-square pixel-wise intensity difference between consecutive greyscale frames and displayed as a 1 s moving RMS envelope. For vCtx MUA spectral analyses during drifting gratings, 3 s windows beginning at stimulus onset were retained only when the entire window occurred within immobility or movement. Mice required ≥ 100 eligible presentations to contribute to either behavioural condition.

### Statistical analysis

No statistical methods were used to predetermine sample sizes. The mouse was the biological replicate for cohort-level inference. Unless otherwise stated, paired mouse-level comparisons were performed using two-sided Wilcoxon signed-rank tests. Regional comparisons of synchronization locking and preferred phase used permutation tests, and circular phase differences between unit classes were assessed using circular median tests. *P* values were Holm-corrected within the comparison families indicated in the corresponding figure legends. Confidence intervals were estimated by bootstrap or, where observations were nested within mice, hierarchical bootstrap by resampling mice and then observations within each selected mouse. Statistical significance was defined as *P* < 0.05 after any stated correction. Circular means and circular median tests were calculated using CircStat for MATLAB.^32^ Analyses were performed using MATLAB R2023b and Python 3.10.

## Use of generative AI

ChatGPT (OpenAI) was used to assist with code development and to provide feedback on manuscript writing. All generated code and suggestions were reviewed and verified by the authors.

## Data Availability

The data analysed in this study are publicly available through the Allen Brain Observatory Visual Coding Neuropixels dataset from the Allen Institute for Brain Science and were accessed using the AllenSDK.

## Code Availability

Custom analysis code will be made available in a public repository upon publication.

## Acknowledgements

We thank F. Helmchen and members of the Helmchen laboratory for discussions and/or comments on the manuscript.

## Author contributions

T.L. and X.Z. conceived the project. T.L. analysed the data and wrote the paper, with X.Z. providing intellectual guidance and input. Both authors revised the manuscript.

## Extended Data Figures

**Extended Data Fig. 1.**
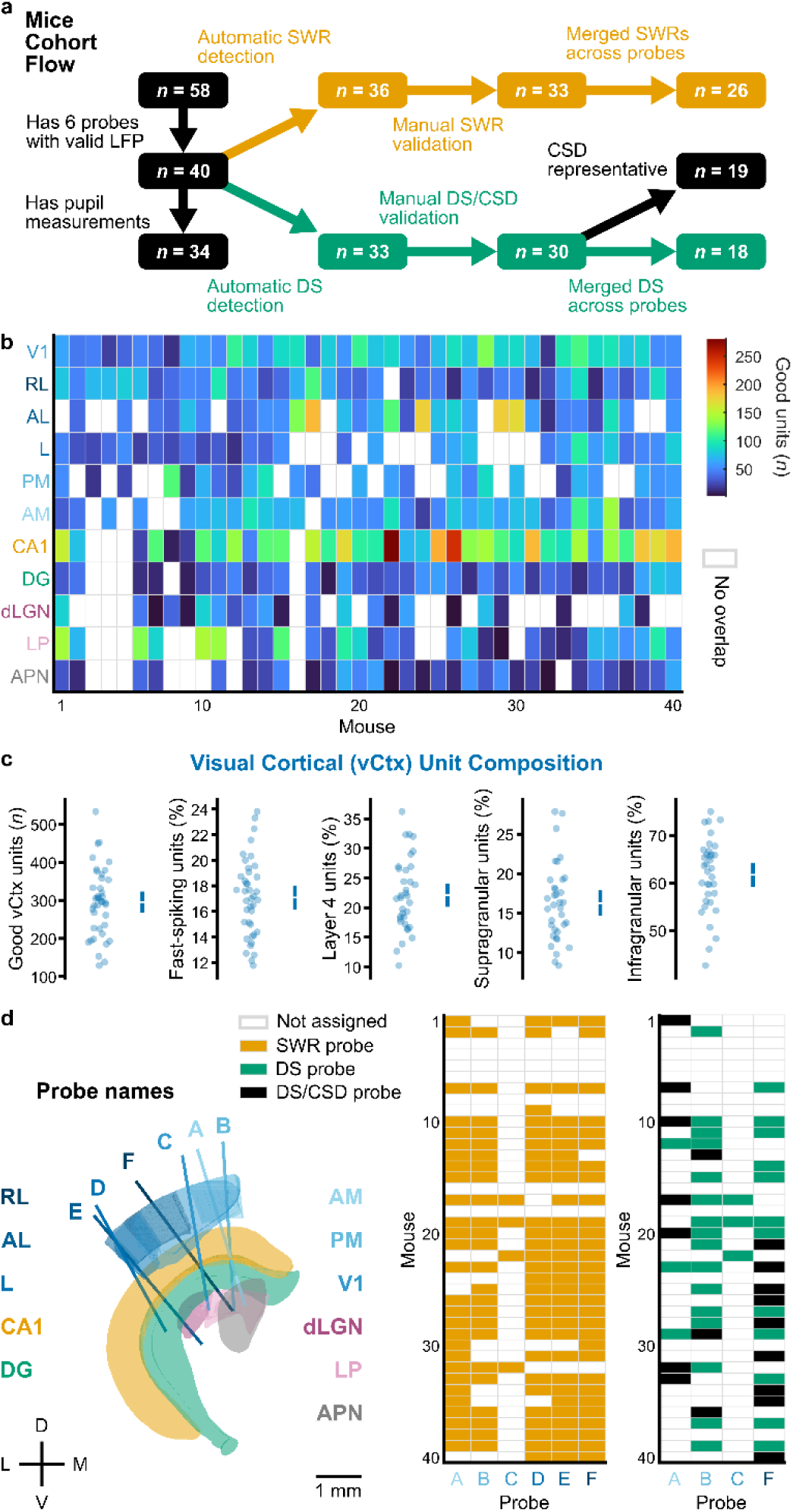
Cohort composition and anatomical sampling. **a**, Cohort selection flow from the Allen Visual Coding Neuropixels dataset to the selected mice for the present analyses. **b**, Number of good units sampled in brainwide areas across included mice for analysis. **c**, Good vCtx unit composition across the cohort. Mice contributed a mean of 295.4 good vCtx units (95% bootstrap CI, 268.1–323.2; *n* = 40 mice), of which 17.08% were putative fast-spiking units (16.17–18.01%; *n* = 40 mice). Among mice with cortical layer assignments (*n* = 35 mice), 22.08% of good vCtx units were in layer 4 (20.12–24.10%), 16.23% were supragranular (14.68–17.87%), and 61.69% were infragranular (59.10–64.17%). Points indicate mice; bars indicate mean and 95% bootstrap CI. **d**, Mean probe trajectories relative to atlas regions (left),^28^ manually validated SWR probes (middle), and manually validated DS probes with probes selected for laminar CSD analysis (right).

**Extended Data Fig. 2.**
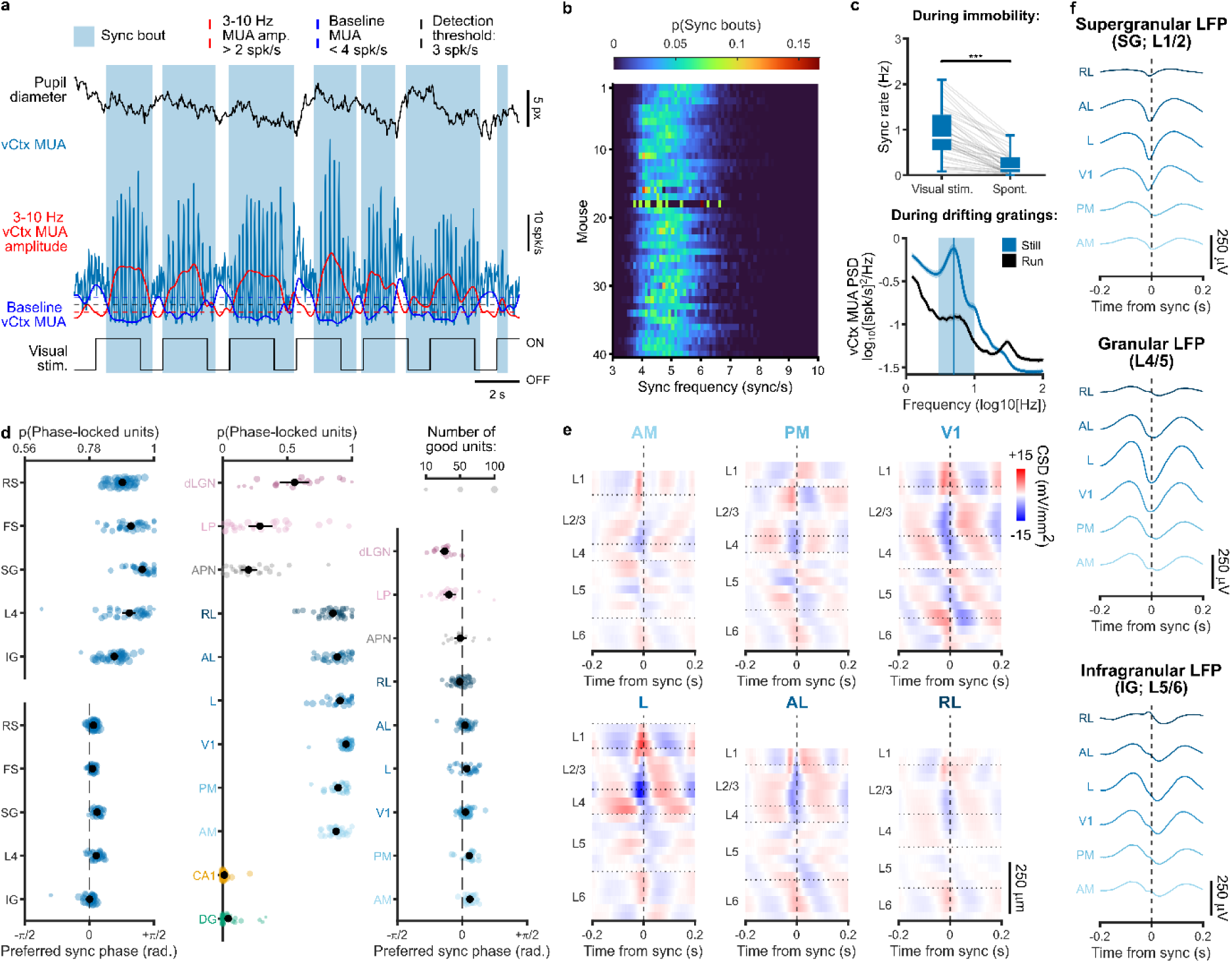
Characterization of cortical synchronization. **a**, Example synchronization bout detection showing aligned pupil diameter, instantaneous vCtx MUA, smoothed 3–10 Hz vCtx MUA amplitude, baseline vCtx MUA, and drifting grating stimulation. Horizontal lines show thresholds for synchronization bout detection. **b**, Normalized synchronization bout frequency distribution across mice. **c**, Synchronization rate during immobile visual stimulation compared to immobile spontaneous activity (*n* = 39 mice; *P* = 5.26 × 10^-8^) (top). vCtx MUA power spectral density during drifting grating stimulation when immobile or moving (bottom). Three second drifting grating trial windows were required to fall entirely within immobility or movement periods, and mice contribute to a behavioural state when at least 100 valid presentations are available. Line and shading indicate mean ± s.e.m., and shaded window and vertical line show 3–10 Hz and 5 Hz, respectively. **d**, Locking of visual cortical units to synchronization phase (left). Fraction of phase-locked units and preferred synchronization phase of regular-spiking (RS), fast-spiking (FS), supragranular (SG), layer 4 (L4), and infragranular (IG) visual cortical units. For the fraction of phase-locked units: RS versus FS/ SG versus L4/ SG versus IG/ L4 versus IG, *n* = 40/ 34/ 34/ 36 paired mice and *P* = 0.004/ 4.09 × 10^-4^/ 1.91 × 10^-6^/ 5.03 × 10^-4^. For preferred phase, corresponding *P* = 0.308/ 0.864/ 1.54 × 10^-4^/ 9.38 × 10^-4^. Similarly, the fraction of phase-locked units and preferred synchronization phase of brainwide units (right). **e**, Example synchronization-aligned CSD across RL, AL, L, V1, PM, and AM. **f**, Synchronization-aligned mean L1/2, L4/5, and L5/6 LFP across RL, AL, L, V1, PM, and AM. For **d**, *P* values were Holm-corrected across the four locking-fraction and four preferred-phase comparisons. \**P* < 0.05, \*\**P* < 0.01, \*\*\**P* < 0.001.

**Extended Data Fig. 3.**
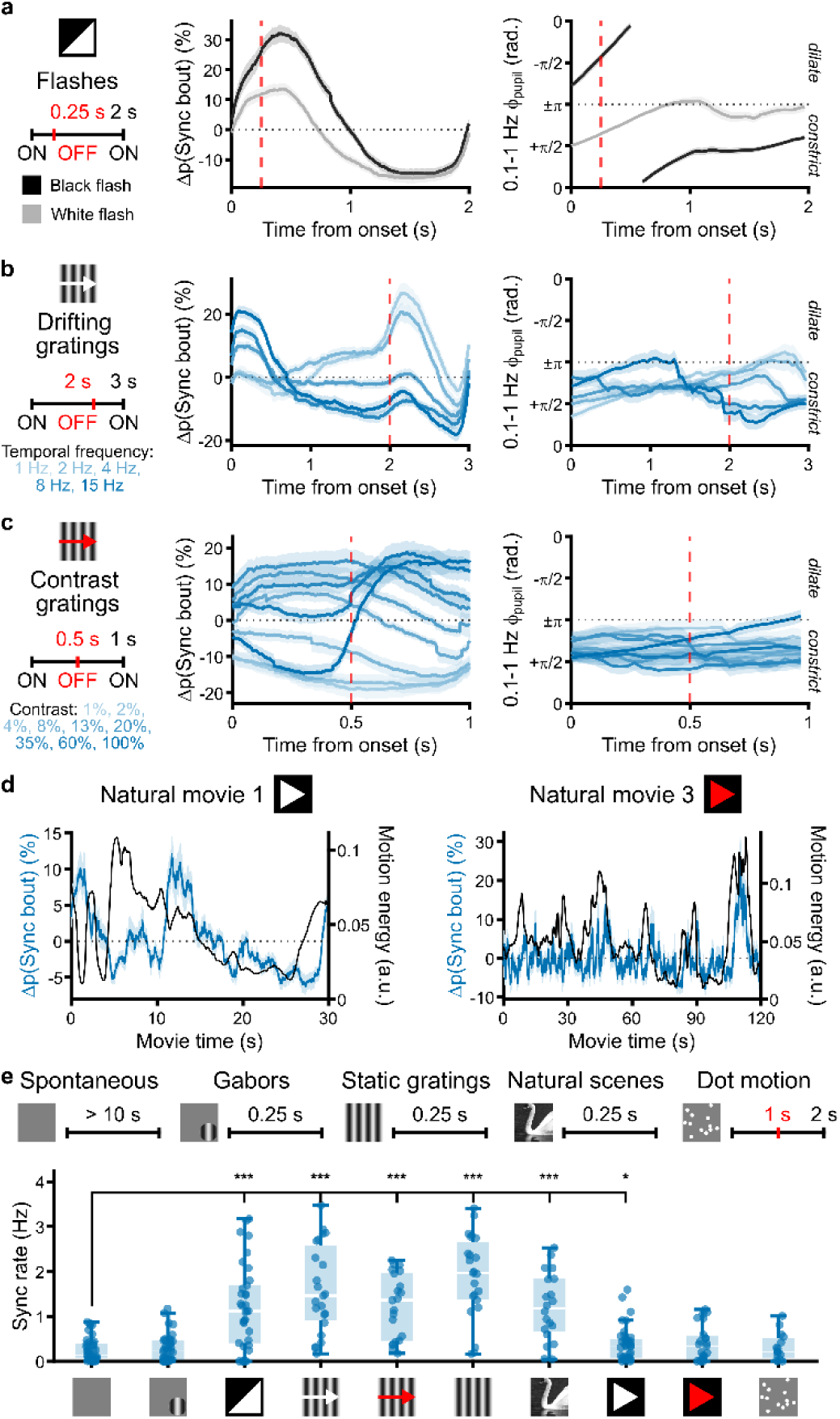
Visual stimulation modulates cortical synchronization. **a**, **b**, **c**, Mean mouse-centered change in synchronization bout likelihood (middle) and mean 0.1–1 Hz pupil phase (right) during full-field flashes (**a**), drifting gratings (**b**), and contrast gratings (**c**). Lines and shading indicate mean ± s.e.m. across mice. **d**, Mean mouse-centered change in synchronization bout likelihood shown together with the smoothed movie motion energy for natural movie 1 (left) and natural movie 3 (right). Synchronization traces show mean ± s.e.m. across mice. **e**, Synchronization rates during various visual stimuli compared with spontaneous activity during immobility. For Gabors/ flashes/ drifting gratings/ contrast gratings/ static gratings/ natural scenes/ natural movie 1/ natural movie 3/ dot motion, *n* = 37/ 34/ 23/ 20/ 23/ 20/ 39/ 23/ 15 mice and *P* = 0.213/ 2.21 × 10^-5^/ 2.16 × 10^-4^/ 5.31 × 10^-4^/ 2.16 × 10^-4^/ 5.31 × 10^-4^/ 0.022/ 0.099/ 0.065. Schematics indicate each stimulus category and its associated trial (black) and offset (red) durations. *P* values were Holm-corrected within each comparison family. \**P* < 0.05, \*\**P* < 0.01, \*\*\**P* < 0.001.

**Extended Data Fig. 4.**
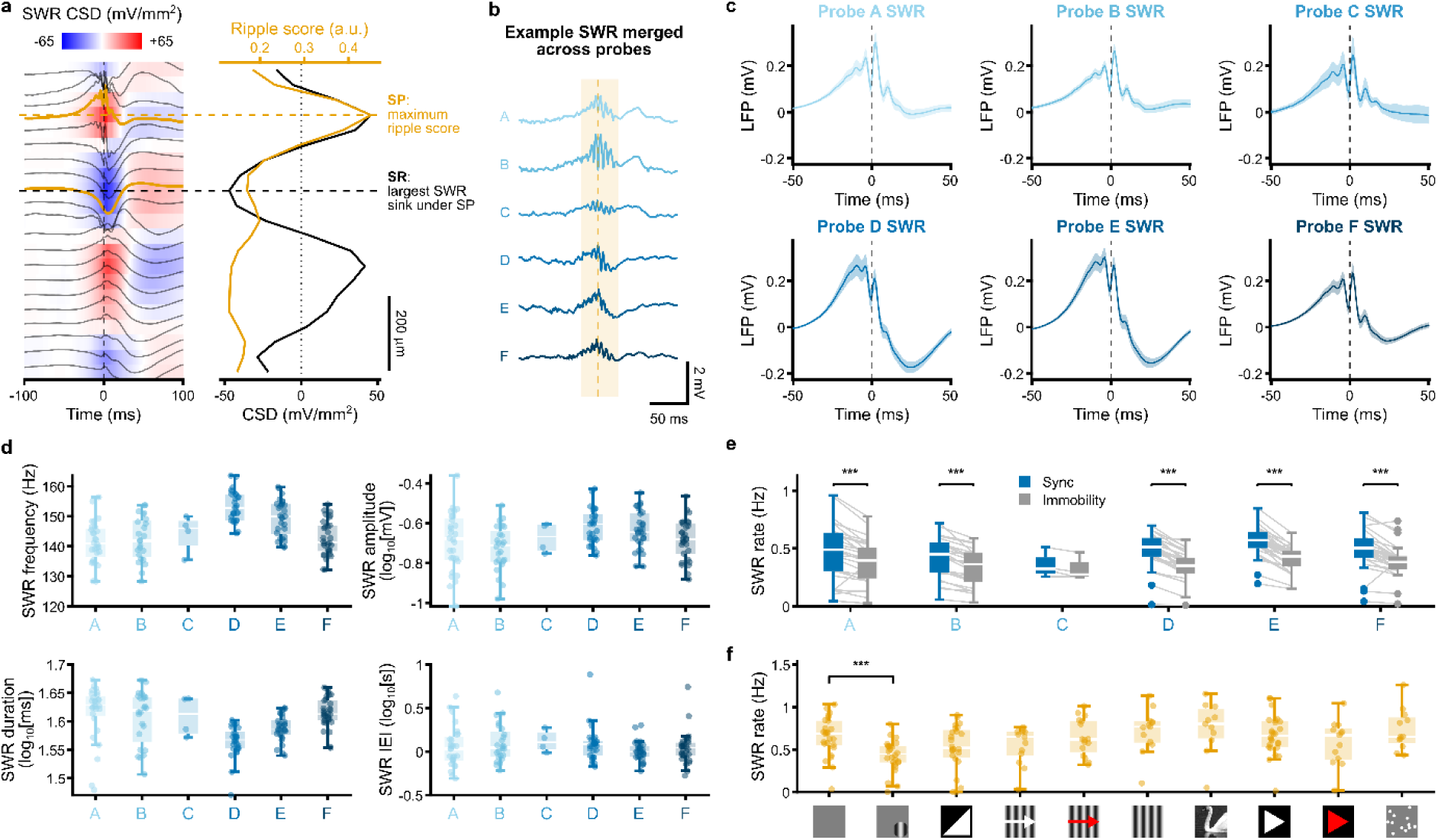
Characterization of sharp-wave ripples. **a**, Example SWR-aligned CA1 CSD and LFP (left) and CSD and ripple score profiles (right) across CA1 channels. **b**, Example SWR merged across 6 probes. Shading indicates the merged SWR window and vertical line the SWR peak. **c**, Probe-wise mean SP LFP waveform aligned to SWR peaks. Lines and shading indicate the mean and 95% hierarchical-bootstrap CI. **d**, Mouse-level SWR frequency, amplitude, duration, and inter-event interval (IEI) across probe range A–F. Amplitude, duration, and IEI are summarized within mouse by geometric means. **e**, SWR rates during synchronization bouts compared with all valid immobility. For probes A/ B/ C/ D/ E/ F, *n* = 28/ 24/ 4/ 29/ 29/ 30 mice, *P* = 1.97 × 10^-4^/ 8.28 × 10^-5^/ 0.250/ 1.54 × 10^-5^/ 1.54 × 10^-5^/ 1.22 × 10^-4^. **f**, SWR rates during various visual stimuli compared with spontaneous activity during immobility. For Gabors/ flashes/ drifting gratings/ contrast gratings/ static gratings/ natural scenes/ natural movie 1/ natural movie 3/ dot motion, *n* = 26/ 24/ 13/ 16/ 13/ 11/ 26/ 13/ 12 mice. Only Gabors differed significantly from spontaneous activity (*P* = 2.65 × 10^-4^; all other adjusted *P* ≥ 0.148). *P* values were Holm-corrected within each comparison family. \**P* < 0.05, \*\**P* < 0.01, \*\*\**P* < 0.001.

**Extended Data Fig. 5.**
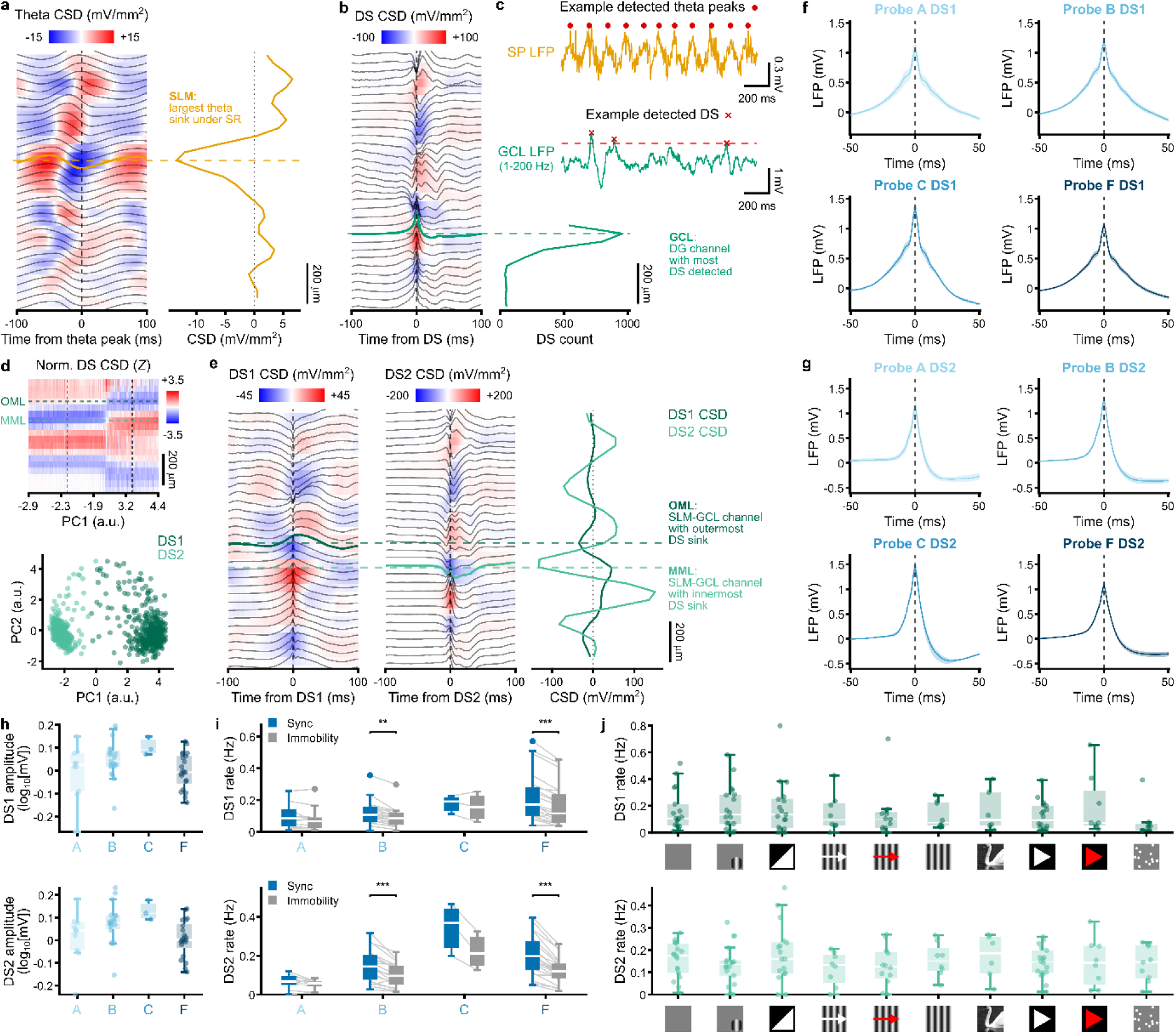
Characterization of dentate spikes. **a**, Example theta-aligned CSD and LFP (left) and CSD profile (right). **b,** Example DS-aligned CSD and LFP (left) and DS counts across DG channels (right). **c**, Example SP and filtered GCL LFP with detected theta peaks and DSs. **d**, Example normalized DS CSD profiles sorted by PC1 (top) and corresponding PC1–PC2 scores classified as DS1 or DS2 (bottom). **e**, Example DS1- and DS2-aligned CSD and LFP (left and middle) and corresponding CSD profiles (right). **f**, **g**, Probe-wise GCL LFP waveforms aligned to DS1 (**f**) and DS2 (**g**). Lines and shading indicate mean and 95% hierarchical-bootstrap CI. **h**, Mouse-level geometric mean of DS1 (top) and DS2 (bottom) amplitudes across probes A, B, C, and F. **i**, DS1 (top) and DS2 (bottom) rates during synchronization bouts compared with immobility epochs. For probes A/ B/ C/ F, DS1: *n* = 10/ 19/ 3/ 21 mice, *P* = 0.320/ 0.002/ 0.500/ 4.91 × 10^-4^; DS2, *n* = 10/ 19/ 3/ 21 mice, *P* = 0.168/ 8.78 × 10^-4^/ 0.250/ 2.38 × 10^-4^. **j**, DS1 (top) and DS2 (bottom) rates during various visual stimuli compared with spontaneous activity during immobility. For Gabors/ flashes/ drifting gratings/ contrast gratings/ static gratings/ natural scenes/ natural movie 1/ natural movie 3/ dot motion, *n* = 18/ 17/ 8/ 12/ 8/ 6/ 18/ 8/ 9 mice. For both DS1 and DS2, no comparison was significant (all adjusted *P* ≥ 0.352). *P* values were Holm-corrected within each comparison family. \**P* < 0.05, \*\**P* < 0.01, \*\*\**P* < 0.001.

**Extended Data Fig. 6.**
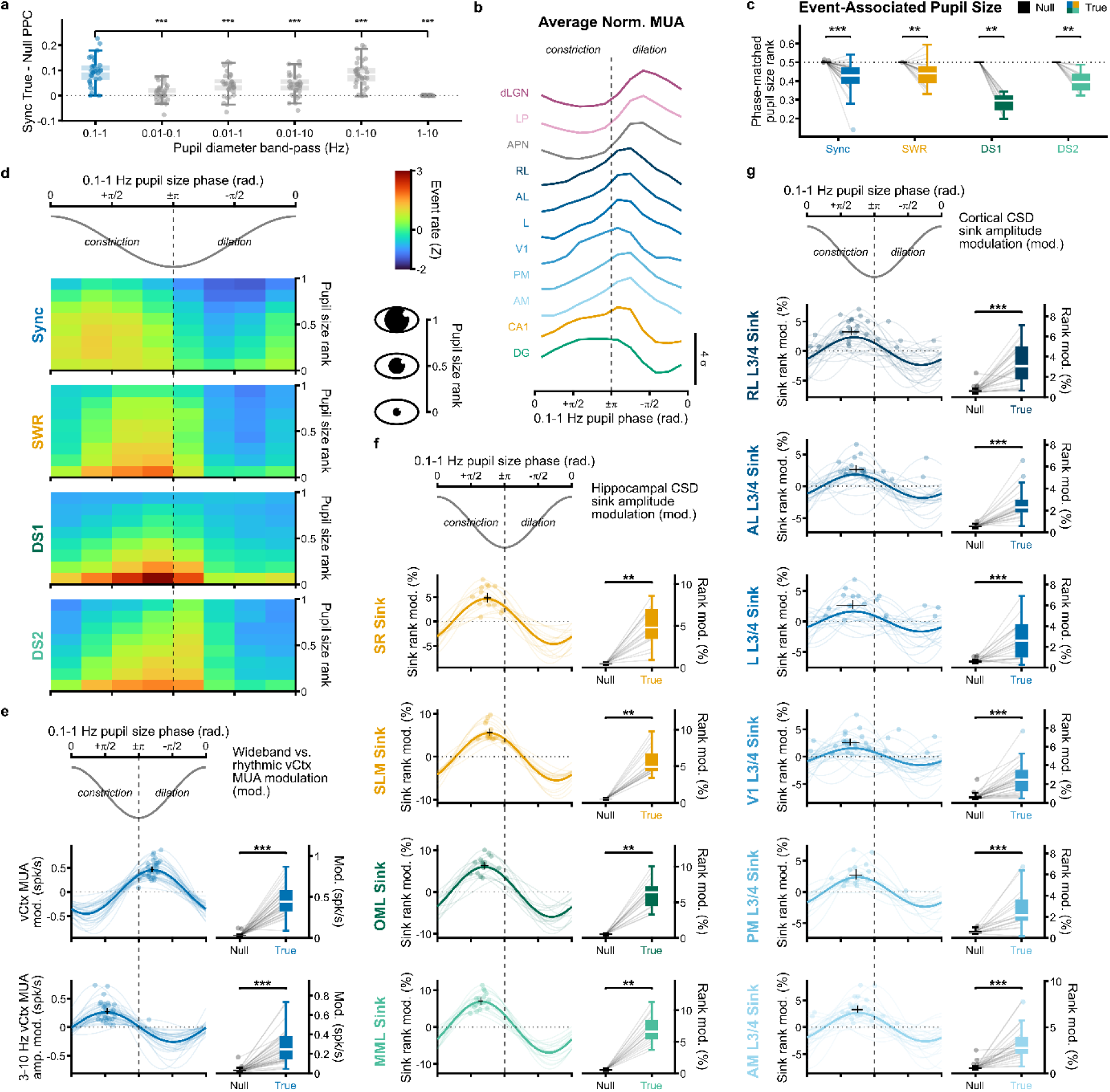
Slow pupil size dynamics during immobility are associated with cortical and hippocampal activity. **a**, Synchronization locking in the 0.1–1 Hz band of pupil size fluctuations was compared with 0.01–0.1/ 0.01–1/ 0.01–10/ 0.1–10/ 1–10 Hz (*n* = 32 mice; P = 8.94 × 10^-6^/ 1.27 × 10^-5^/ 1.27 × 10^-5^/ 1.27 × 10^-5^/ 4.81 × 10^-6^). **b**, Average brainwide MUA profiles across 0.1–1 Hz pupil phase during spontaneous immobility. **c**, Pupil-size ranks of synchronization, SWR, DS1, and DS2 relative to phase-matched pupil-cycle-shift null ranks (Sync/ SWR/ DS1/ DS2, *n* = 34/ 22/ 16/ 16 mice; *P* = 5.96 × 10^-6^/ 0.001/ 0.001/ 0.001). **d**, Mean *Z*-scored synchronization, SWR, DS1, and DS2 rate as a function of pupil phase and pupil size rank. **e**, Modulation of vCtx MUA and 3–10 Hz MUA amplitude across pupil phase during immobility (*n* = 34 mice; *P* = 7.31 × 10^-7^/ 7.31 × 10^-7^). **f**, Modulation of discrete SR, SLM, OML, and MML CSD sink strength rank across pupil phase during spontaneous immobility (*n* = 17 mice; *P* = 0.001/ 0.001/ 0.001/ 0.001). **g**, Modulation of discrete vCtx L3/4 CSD sink strength across pupil phase during spontaneous immobility (RL/ AL/ L/ V1/ PM/ AM, *n* = 27/ 24/ 21/ 29/ 17/ 27 mice; *P* = 4.4 × 10^-5^/ 6.21 × 10^-5^/ 3.73 × 10^-4^/ 1.71 × 10^-5^/ 3.73 × 10^-4^/ 2.8 × 10^-5^). For **e**–**g**, left plots show mouse-level fitted modulation and the cohort mean; crosses indicate cohort preferred phase and modulation with 95% hierarchical-bootstrap CIs, and right plots compare true and null modulation. *P* values were Holm-corrected within each comparison family. \**P* < 0.05, \*\**P* < 0.01, \*\*\**P* < 0.001.

**Extended Data Fig. 7.**
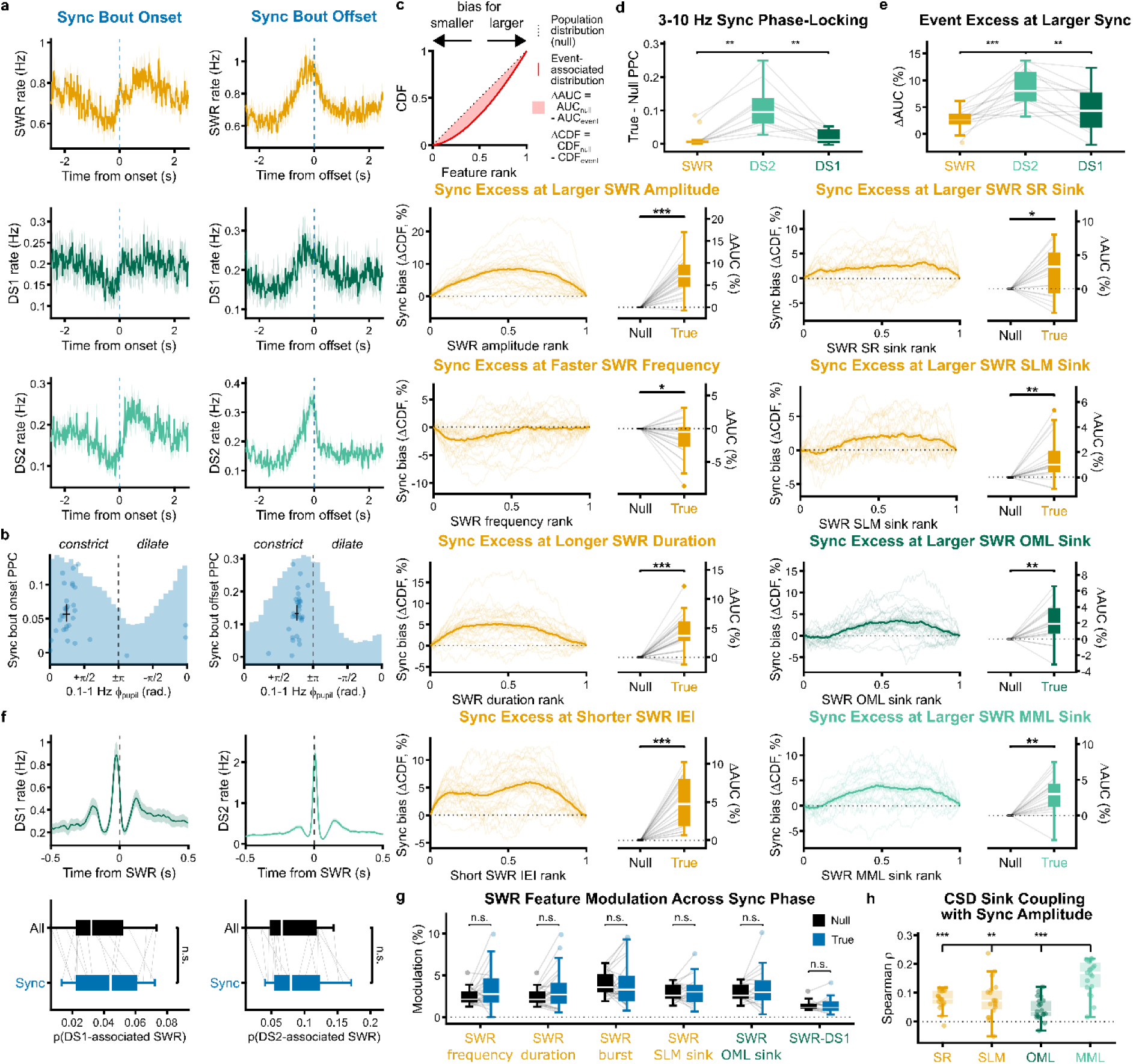
Cortical synchronization is associated with the timing and properties of hippocampal events. **a**, SWR, DS1, and DS2 rates aligned to synchronization bout onset (left) and offset (right) (SWR, *n* = 26 mice; DS1/DS2, *n* = 18 mice). Lines and shading indicate mean ± s.e.m. across mice. **b**, Locking of synchronization bout onset and offset to 0.1–1 Hz pupil size phase. Crosses indicate preferred phase and PPC with 95% hierarchical-bootstrap CI across mice. Observed PPC was compared with circular-shift null PPC within mouse (onset: *n* = 31 mice, *P* = 3.1 × 10^-6^; offset: *n* = 31 mice, *P* = 2.59 × 10^-6^). **c**, Bias of synchronization-associated SWR across intrinsic SWR feature and laminar sink ranks. Curves show synchronization-associated rank bias (ΔCDF; left), and boxplots ΔAUC (AUC − 50%; right). Intrinsic features, *n* = 26 mice: amplitude/ frequency/ duration/ short IEI, *P* = 3.32 × 10^-5^/ 0.041/ 4.2 × 10^-5^/ 3.32 × 10^-5^. Sinks, *n* = 19 mice: SR/ SLM/ OML/ MML, *P* = 0.010/ 0.004/ 0.007/ 0.002. **d**, **e**, Matched SWR, DS2, and DS1 synchronization phase-locking (**d**; *n* = 11 mice; SWR–DS1/ SWR–DS2/ DS1–DS2, *P* = 0.577/ 0.003/ 0.004) and bias toward stronger synchronized bursts (**e**; *n* = 15 mice; *P* = 0.073/ 3.66 × 10^-4^/ 0.002). **f**, DS1 and DS2 rates around SWR peaks and proportion of SWRs associated with DS1 or DS2 among all versus synchronization-associated SWRs (*n* = 17 mice; DS1/ DS2, *P* = 0.113/ 0.099). **g**, Additional modulation across synchronization phase versus permutation nulls (frequency/ duration/ burst/ SLM/ OML/ DS1, *n* = 26/ 26/ 26/ 19/ 19/ 15 mice; *P* = 0.138/ 0.276/ 1/ 1/ 1/ 1). **h**, Coupling between synchronization amplitude and hippocampal CSD sink strength (*n* = 19 mice). MML versus SR/ SLM/ OML, *P* = 9.27 × 10^-4^/ 0.002/ 8.78 × 10^-4^. *P* values were Holm-corrected within each comparison family. \**P* < 0.05, *\*\*P* < 0.01, \*\*\**P* < 0.001.

